# Retinoic acid organizes the vagus motor topographic map via spatiotemporal regulation of Hgf/Met signaling

**DOI:** 10.1101/826735

**Authors:** Adam J. Isabella, Gabrielle R. Barsh, Jason A. Stonick, Cecilia B. Moens

## Abstract

The topographic map, in which the positions of neuron cell bodies correspond with the positions of their synaptic targets, is a major organizational motif in the nervous system. To understand how topographic axon targeting is controlled during development, we examine the mechanism underlying topographic innervation of the pharyngeal arches by the vagus motor nerve in zebrafish. We reveal that Retinoic Acid organizes the topographic map by specifying anterior-posterior identity in post-mitotic vagus motor neurons. We then show that chemoattractant signaling between hepatocyte growth factor (Hgf) and the Met receptor is required for pharyngeal arch innervation by the vagus motor nerve. Finally, we find that Retinoic Acid controls the spatiotemporal dynamics of Hgf/Met signaling to coordinate axon targeting with the developmental progression of the pharyngeal arches and show that experimentally altering the timing of Hgf/Met signaling is sufficient to redirect axon targeting and disrupt the topographic map. These findings establish a new mechanism of topographic map development in which regulation of chemoattractant signaling in both space and time guides axon targeting.

## INTRODUCTION

In order for information to be accurately transmitted through the complex cellular networks that comprise the nervous system, the pattern of connectivity within these networks must be extremely well organized. Many neural networks are organized into topographic maps, a spatial patterning principle in which the relative positions of neurons’ cell bodies in a projecting field corresponds to the relative positions of their axons in the target field. Topographic organization maintains the spatial component of sensory or motor information during transmission, and is believed to be crucial for coherent, accurate, and efficient flow of information through the complex neural circuitry (Bednar, 2016; Cang and Feldheim, 2013; Chklovskii and Koulakov, 2004; Levine et al., 2012). Disruption of these maps can impair signal coordination and processing and, ultimately, behavior (Bargary and Mitchell, 2008; Sperry, 1943; Sürmeli et al., 2011). Although most, if not all, sensory and motor systems are topographically organized, in many cases we do not understand how these maps form during embryonic development.

Topographic innervation patterns are established by the guidance of growing axons to the appropriate target regions during development. The predominant model of topographic mapping is one of spatial signaling, in which growing axons interpret spatially patterned chemoattractant and/or chemorepulsive cues to reach appropriate targets (Sperry, 1963). A classic example of this model is the retinotopic map, in which retinal neurons project axons to the optic tectum. In this system, corresponding gradients of Ephrin ligands and Eph receptors in the retina and tectum allow retinal axons to select their target site by seeking the optimal Eph-Ephrin signaling state (Triplett and Feldheim, 2012).

While retinal axon targeting relies purely on the conveyance of spatial information, in some cases the timing of developmental events is an important component of topographic map development (Eerdunfu et al., 2017; Kulkarni et al., 2016; Petrovic and Hummel, 2008; Takeuchi et al., 2010). For example, topographic mapping of mouse olfactory sensory neurons is instructed by their sequential projection of axons to the target field (Eerdunfu et al., 2017; Takeuchi et al., 2010). In this and other such cases, temporal differences in neuronal identity and targeting have been ascribed to differences in the timing of neuronal birth. It remains unclear whether and how a temporal mechanism might guide projections of groups of neurons in the brain that share a common birth time. Additionally, it is not clear how changes in molecular communication between neurons and their targets are coordinated over time to ensure precise axon-target matching over the period of axonal projection.

We recently identified a key role for developmental timing in the topographic organization of the zebrafish vagus nerve, which functions independently of neuron birth timing (Barsh et al., 2017). The vagus nerve (cranial nerve X) carries sensory information of multiple modalities from the pharynx, larynx, heart and viscera, and returns appropriate motor and parasympathetic innervation for control of speech, swallowing, heart rate, and multiple aspects of digestion. How vagus sensorimotor circuits are spatially coordinated is not currently understood. In zebrafish, Vagus motor neurons (mX neurons) are born ventrally in the posterior hindbrain, and migrate dorsolaterally to form a discrete nucleus of a few hundred cells between 24 and 36 hours post fertilization (hpf). After reaching the nucleus, mX neurons extend axons that collect in a large fascicle that projects anteriorly towards the otic vesicle and then turns ventrally toward the pharyngeal arch (PA) region (Fig. 1A). Before entering the PAs, the nerve splits into five branches, four of which innervate PAs 4, 5, 6, & 7 (branches 4, 5, 6, & 7), and one of which extends posteriorly to innervate the viscera (visceral branch) (Fig. 1A). Each mX neuron extends an axon into a single branch (Barsh et al., 2017). Innervation occurs between 28 and 60 hpf. The vagus motor nerve exhibits a topographic organization that is conserved between fish and mammals, with more anterior neurons innervating more anterior branches and more posterior neurons innervating more posterior branches (Fig 1B) (Barsh et al., 2017; Bieger and Hopkins, 1987; Morita and Finger, 1985). We found that mX neurons at different A-P positions extend axons and innervate their targets at different developmental times, and that experimentally changing the timing of axon outgrowth in individual mX neurons is sufficient to override spatial cues and alter axon target selection (Barsh et al., 2017). Thus, mX neurons must integrate spatial and temporal cues to coordinate topographic axon targeting decisions. However, we do not know the nature of these cues, their dynamics in space and time, or how they are coordinated between mX neurons and their targets to precisely guide targeting.

**Figure 1:**
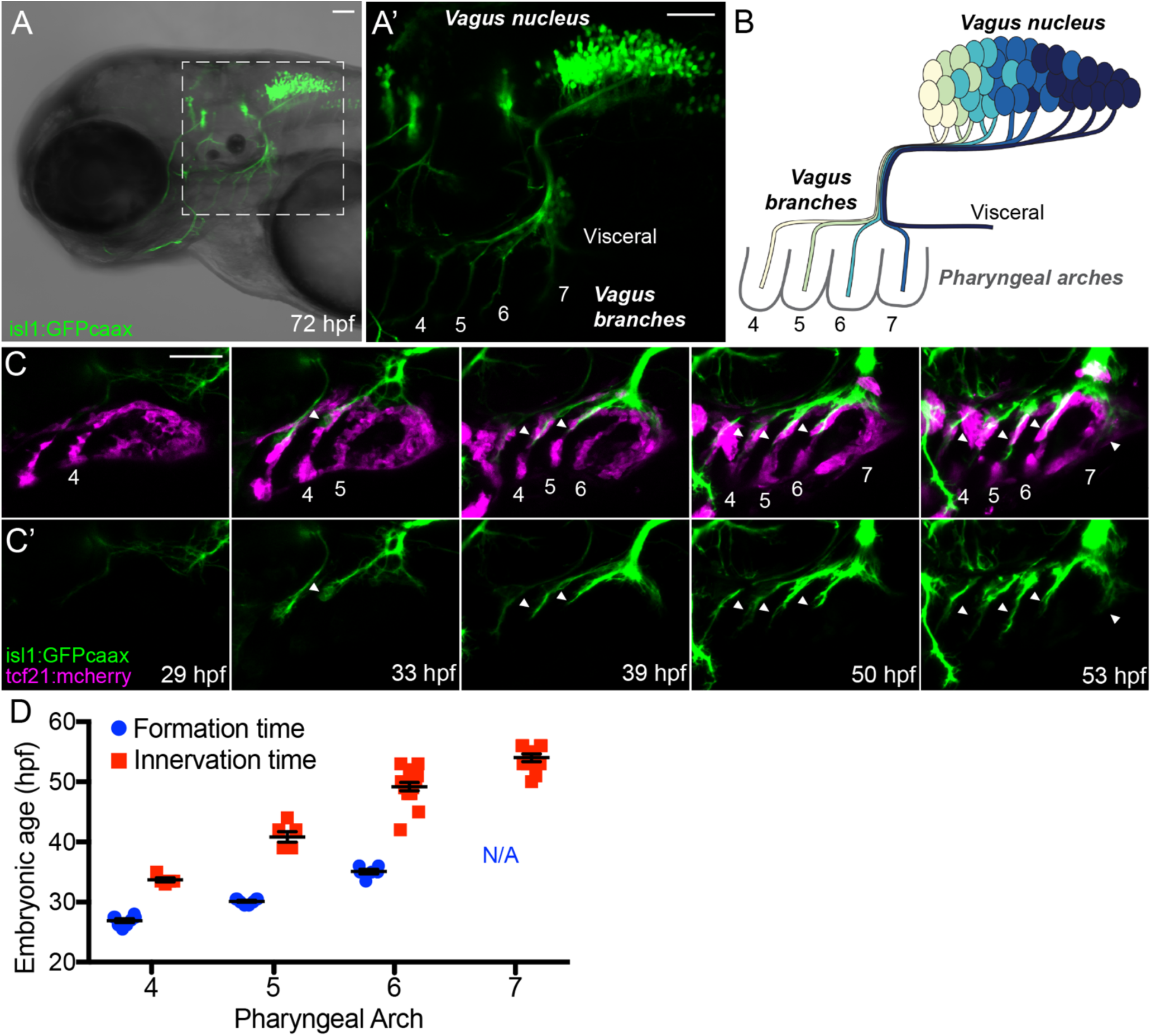
Spatiotemporal dynamics of vagus nerve branch development. (A) Zebrafish vagus nerve structure. (A’) Enlarged view of boxed region. Neurons in the vagus nucleus extend axons into five pharyngeal arch branches (4-7 and Visceral). (B) Vagus nerve topography. Cells in the anterior of the nucleus (yellow and green) innervate anterior branches. Cells further posterior in the nucleus (blues) innervate more posterior branches. (C) Time series of PA4-7 formation and innervation. PAs (marked by tcf21:mCherry expression in muscle progenitors, magenta) form sequentially from anterior to posterior and are sequentially innervated by mX axons (isl1:GFPcaax, green). Numbers indicate PAs. Arrowheads indicate axon branches. (D) Quantification of PA formation and innervation timing represented in (C). Each point represents a single embryo in which the PA indicated was beginning to be innervated. Data represent mean ± SEM. All images are lateral views oriented with anterior to left. Scale bars = 50 μm.

Here, we examine the cell signaling dynamics within and between vagus neurons and the pharyngeal arches that regulate topographic vagus axon targeting. We find that Retinoic Acid (RA) signaling in the posterior hindbrain directs axon target selection by providing A-P positional information to mature mX neurons. We then identify the Met receptor and its ligand, Hgf, as the chemoattractant signaling system that guides mX axon targeting. Finally, we discover that RA drives a spatiotemporal wave of *met* expression in mX neurons that is coordinated with the sequential onset of *hgf* expression in the PAs. By manipulating the dynamics of Hgf/Met signaling, we reveal how the spatiotemporal regulation of chemoattraction controls topographic axon targeting in this system. This work establishes a new model of topographic map development in which neurons and their targets coordinate chemoattractant signaling in both space and time to guide axon targeting.

## RESULTS

### Spatiotemporal dynamics of pharyngeal arch formation and innervation

We previously identified a critical role for innervation timing in the formation of the vagus topographic map (Barsh et al., 2017). In order to elucidate the mechanisms that guide timing-dependent targeting, we first established a precise developmental timeline of the formation of pharyngeal arches 4-7 and their innervation by mX neurons. We collected a carefully staged image series of 24-60 hpf embryos expressing fluorescent markers of the PAs (*TgBAC(tcf21:mcherry-NTR)*, which is expressed in PA muscle precursors) (Nagelberg et al., 2015; Wang et al., 2015), and mX neurons (*Tg(isl1:GFPcaax)*, which is expressed in in post-mitotic cranial motor neurons) (Fig. 1C) (Higashijima et al., 2000). We found that the PAs form by sequentially emerging from a posterior tcf21:mcherry+ primordium with the following timing: PA4: 27 hpf; PA5: 30 hpf; PA6: 35 hpf (Fig. 1C-D). The timing of PA7 formation cannot be determined as it appears to derive from what remains of the primordium after PA6 emerges. We found that mX axons innervate the PAs sequentially with the following timing: PA4: 34 hpf; PA5: 41 hpf; PA6: 49 hpf; PA7: 54 hpf (Fig. 1C-D). These results provide a timeline with which to contextualize the timing of events reported in this study.

### Retinoic Acid is a putative regulator of A-P vagus motor neuron identity

The A-P position of a mX neuron within the vagus nucleus determines which branch it innervates, indicating that mX neurons exhibit molecular identity differences along the A-P axis that control axon targeting (Barsh et al., 2017). Furthermore, experimentally altering the A-P position of a post-mitotic mX neuron is sufficient to change target selection, indicating the presence of inductive signals along the A-P axis of the mX nucleus (Barsh et al., 2017). The signals that regulate A-P mX identity, and how these signals control axon targeting, are not known. We reasoned that understanding the gene expression differences between anterior and posterior mX neurons would provide valuable insight into how they select different axon targets. To identify these differences, we performed bulk RNA sequencing (RNA-seq) on anterior and posterior mX neurons at 28-30 hpf, shortly after the onset of axon formation. To isolate these cell populations, we expressed the photoconvertible protein Kaede, which switches from green to red fluorescence upon illumination with ultraviolet light, in mX neurons (*Tg(isl1:Kaede)*). We then photoconverted either anterior or posterior mX neurons (Fig. 2A), dissected and dissociated the posterior hindbrain to single-cell suspension, and manually collected the photoconverted neurons. We sequenced cDNA isolated from 3 replicates of 100 cells per position. Overall, anterior and posterior mX neurons have very similar transcriptomes; however, we identified several genes enriched in anterior or posterior neurons (Fig. 2B). In validation of this approach, one of the most significantly posteriorly-enriched genes it identified is *hoxb5a*, which we previously found to be expressed only in posterior mX neurons (Fig. 2C) (Barsh et al., 2017).

**Figure 2:**
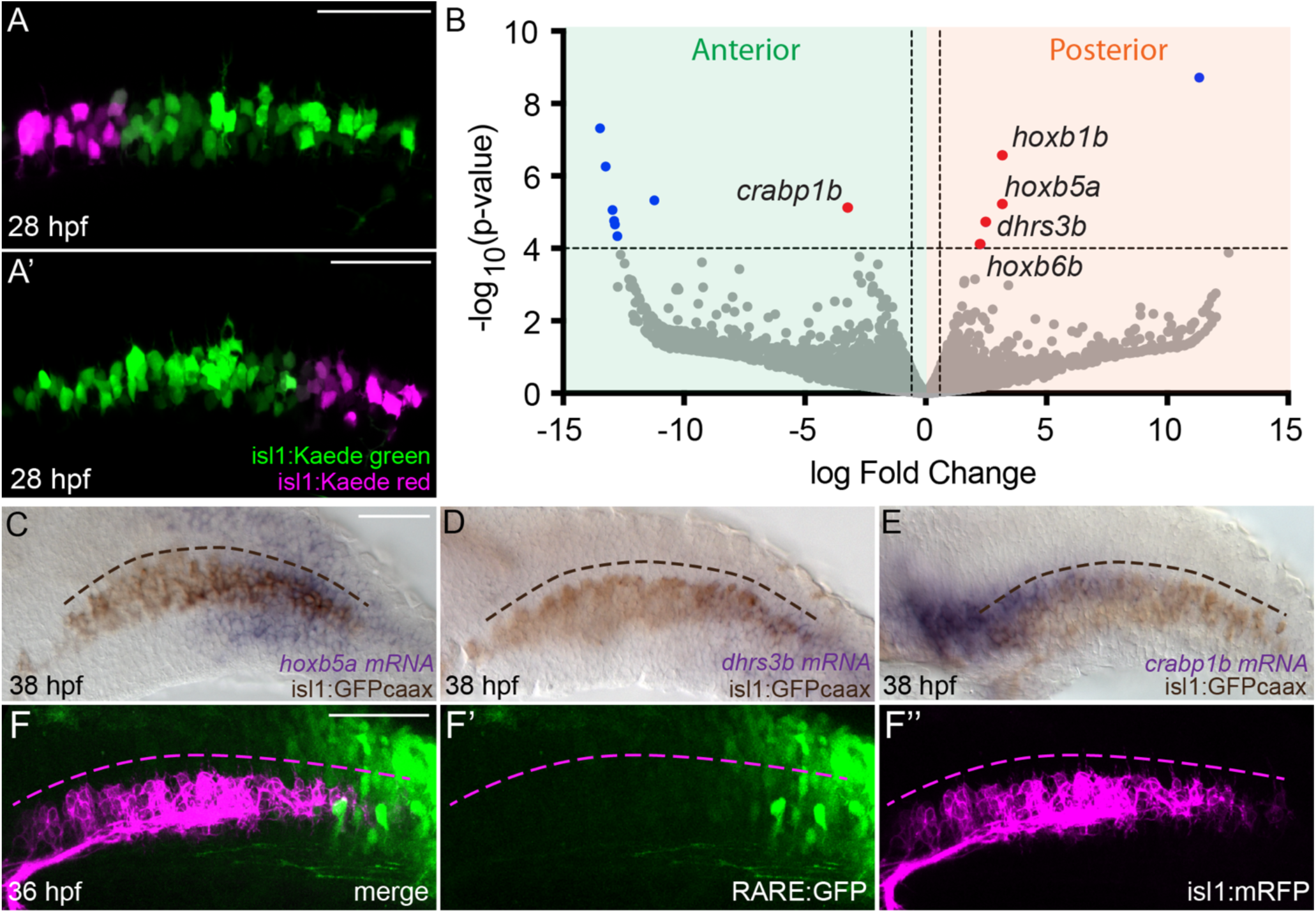
Retinoic Acid is a putative regulator of A-P vagus motor neuron identity. (A-B) Differential gene expression between anterior and posterior mX neurons. (A) Representative anterior (A) and posterior (A’) photoconverted (magenta) regions collected for RNAseq analysis. (B) Volcano plot of RNAseq data indicating mRNAs enriched in anterior (left) or posterior (right) mX neurons. Dashed lines indicate significance threshold for a false discovery rate of 10% (y-axis) and a fold change of 1.5 (x-axis). Blue and red dots represent significantly differentially expressed genes. Red dots represent genes indicative of differential Retinoic Acid signaling between populations. (C-E) RNA *In situ* hybridization of selected genes identified in (B). In each image, mRNA expression is purple and the mX neurons marked by isl1:GFP are brown. The A-P span of the mX nucleus is indicated by the curved dotted line. *hoxb5a* (C) and *dhrs3b* (D) mRNAs are enriched in posterior mX neurons. *crabp1b* (E) mRNA is enriched in anterior mX neurons. (F) The retinoic acid-responsive RARE:GFP transgene (green) is expressed in posterior, but not anterior, mX neurons (magenta). All images are lateral views oriented with anterior to left. Scale bars = 50 μm.

Our results reveal the extrinsic signaling factor that specifies A-P mX neuron identity. Specifically, after filtering out genes with no expression in one population, the five most significantly differentially expressed genes (highlighted in Fig. 2B) are transcriptional targets of Retinoic Acid (RA) signaling (*hoxb1b, hoxb5a, hoxb6b, dhrs3b*, all posteriorly enriched) (Chen et al., 2007; Kam et al., 2013; Kudoh et al., 2002; Oosterveen et al., 2003) and/or regulators of RA signaling (*dhrs3b, crabp1b*) (Cai et al., 2012; Feng et al., 2010; Kam et al., 2013). We validated the differential expression of *dhrs3b* and *crabp1b* with RNA *in situ* hybridization (ISH) (Fig. 2D-E). These data suggest that posterior mX neurons experience higher levels of RA than anterior mX neurons.

RA is a morphogen that controls A-P embryonic patterning, primarily through the transcriptional regulation of *Hox* genes (Nolte et al., 2019; Rhinn and Dolle, 2012). RA is produced in the paraxial mesoderm posterior to the hindbrain, and plays a crucial early (before 14 hpf) role in the A-P patterning of the hindbrain (Moens and Prince, 2002; Nolte et al., 2019), as well as in the patterning of spinal motor neurons (Diez del Corral et al., 2003; Novitch et al., 2003). Although RA can play a late (post-24 hpf) role in nervous system patterning in some contexts, such as in guiding enteric neural crest migration (Uribe et al., 2017), it is believed that its ability to specify A-P identity in the hindbrain is restricted to a critical window early in development (Begemann and Meyer, 2001). It is therefore unclear whether RA can influence the anterior-posterior identity of mature vagus neurons. To address this question, we first determined the state of RA signaling in the mX nucleus. We directly visualized RA signaling using a *12xRARE:GFP* transgene, in which GFP is under the control of 12 concatenated Retinoic Acid Response Elements (RAREs) – gene regulatory elements that are bound by Retinoic Acid Receptors (RARs) and mediate RA-dependent gene expression in a dose-dependent manner (Mark et al., 2004; Waxman and Yelon, 2011). Consistent with our RNA-seq data, we observed that *RARE:GFP* signal is high in the posterior and low in the anterior of the vagus nucleus (Fig. 2F). It is important to note that, due to differing sensitivities to RA, individual RA-responsive genes may have expression boundaries that lie anterior or posterior to the *RARE:GFP* expression boundary (Papalopulu et al., 1991). Regardless, the pattern of RA-responsive gene expression we observe across the mX nucleus is consistent with a role for RA in specifying A-P mX neuron identity.

### Retinoic Acid controls axon target selection in mature vagus motor neurons

We hypothesize that RA specifies an aspect of posterior mX neuron identity which promotes targeting to posterior branches and/or prevents targeting to anterior branches. While it is present at the right time and place to play this role, whether RA controls target selection during the period of mX axon outgrowth is unknown. We therefore sought to test whether altering RA signaling levels can change mX axon target selection. We first examined mX axon targeting in *cyp26a1^rw716^* mutant embryos, which are deficient in RA degradation and consequently experience an anterior shift in RA signaling (Emoto et al., 2005). *cyp26a1^rw716^* embryos do not exhibit gross defects in vagus motor nerve structure; however, we predicted that the increased RA levels experienced by anterior neurons in this condition would bias their axons towards more posterior branches. To address this hypothesis, we photoconverted the anterior-most 10% of mX neuron soma in wild-type and *cyp26a1^rw716^ Tg(isl1:Kaede)-*expressing embryos and, after allowing for anterograde transport of the photoconverted protein, we quantified the distribution of photoconverted axons by measuring the red fluorescence signal (normalized to total fluorescence) in each PA branch (Fig. 3A-B). We observed a decrease in the normalized red fluorescence in PA4, indicating that the proportion of anterior mX neurons innervating this branch in *cyp26a1^rw716^* mutants is decreased, and an increase in the normalized red fluorescence in PA7, suggesting that some of the anterior mX neurons that fail to target PA4 are redirected to PA7 in these mutants (Fig. 3B). These data indicate that RA promotes targeting of mX axons to posterior branches.

**Figure 3:**
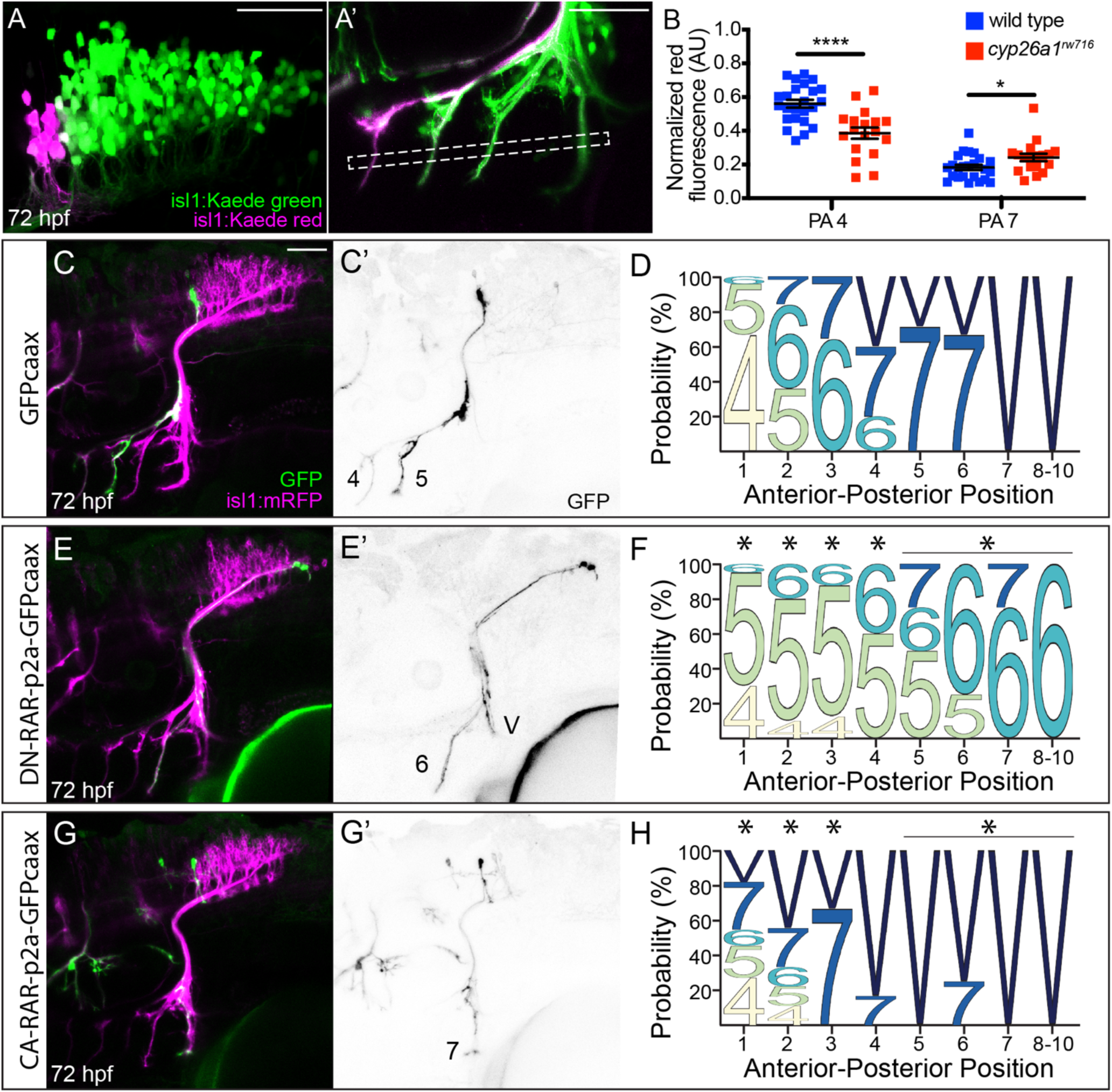
Retinoic Acid controls axon target selection in vagus motor neurons. (A-B) Elevated RA levels in *cyp26a1^rw716^* mutants bias targeting of anterior mX neurons to posterior branches. (A) Example of photoconverted anterior neurons (magenta) (A) and their axons (A’). White box in (A’) shows region used for measurement of axon branch fluorescence intensity. (B) Normalized Kaede-red branch fluorescence intensities in anterior (PA4) and posterior (PA7) branches. Decreased PA4 red signal and increased PA7 red signal indicate that the axons of anterior mX neurons are biased towards more posterior branches in *cyp26a1^rw716^* mutants. Data represent mean ± SEM. t-test ****P<0.0001, *P<0.05. (C-H) Altered RA signaling levels cell-autonomously alters axon target selection in mX neurons. (C) Anterior neurons expressing control *Tg(UAS:GFPcaax)* (green in C, black in C’) innervate anterior branches (PA4, PA5). (D) Position probability matrix showing wild-type vagus topography. N = 89 cells. (E) Posterior neurons expressing *Tg(UAS:DN-RAR-p2a-GFPcaax)* (green in E, black in E’) aberrantly targeting a more anterior branch (PA6). (F) Position probability matrix showing an anterior targeting bias in DN-RAR vagus topography. N = 67 cells. (G) An anterior neuron expressing *Tg(UAS:CA-RAR-p2a-GFPcaax)* (green in G, black in G’) aberrantly targeting a more posterior branch (PA7). (H) Position probability matrix showing a posterior targeting bias in CA-RAR vagus topography. N = 41 cells. (D, F, H) Data are represented as the proportion of cells that innervate the noted branch. Fisher’s exact test comparing positional axon distribution to control, *P<0.05. All images are lateral views oriented with anterior to left. Scale bars = 50 μm.

While the above experiment offers strong evidence that RA regulates mX axon targeting, the constitutive nature of the *cyp26a1* mutation does not allow us to determine if the effect results directly from changes in RA signaling in the motor neurons or indirectly due to changes in other tissues, nor does it indicate when in development this effect occurs. We therefore devised an experiment that tests whether directly altering the levels of RA signaling in mature mX neurons is sufficient to impact axon target selection. For this experiment, we manipulated RA signaling levels in one or a few post-mitotic mX neurons via mosaic expression of UAS-driven transgenes using an *Tg(isl1:Gal4)^fh452^* driver (Davey et al., 2016). Three transgenic constructs were expressed in this manner: 1. *Tg(UAS-GFPcaax)*, which expresses a membrane GFP; 2. *Tg(UAS:DN-hRARα-p2a-GFPcaax),* which expresses a dominant-negative RAR (DN-RAR) in combination with membrane GFP; 3. *Tg(UAS:CA-RARga-p2a-GFPcaax*), which expresses a constitutively active RAR (CA-RAR) in combination with membrane-GFP. The *DN-hRARα* and *CA-RARga* transgenes cell-autonomously decrease and increase RA signaling levels, respectively, in zebrafish (Waxman and Yelon, 2011). The RAR and GFPcaax genes were co-expressed on a single transcript separated by a p2a self-cleaving peptide to ensure that the GFP signal accurately reflects RAR expression (Kim et al., 2011). We then quantified both the A-P position of each GFP+ cell body within the mX nucleus (marked with *Tg(isl1:mRFP*)), and the branch innervated by its axon, to precisely measure the effect of RA signaling levels on individual axon targeting decisions (Fig. 3C-H). This experiment revealed a dramatic cell-autonomous influence of RA signaling on mX axon target selection. Decreasing signaling with *DN-hRARα* causes mX neurons to project to more anterior PAs than is appropriate for their A-P position (Fig. 3E-F). Conversely, increasing RA signaling with *CA-RARga* causes mX neurons to project to more posterior PAs than is appropriate for their A-P position (Fig 3G-H). These data confirm that RA acts to control the axon targeting decisions of mature neurons in the mX nucleus, with low RA levels promoting anterior target selection and high RA levels promoting posterior target selection.

### Retinoic Acid controls mX chemoattraction through Hgf/Met signaling

How does RA control axon target selection? We previously reported that the timing of axon initiation is an important consequence of mX neuron positional identity – anterior mX neurons form axons several hours before posterior mX neurons, and delaying axon outgrowth is sufficient to misdirect the axons of anterior mX neurons to more posterior targets (Barsh et al., 2017). A simple hypothesis, therefore, would be that RA promotes the posterior targeting of posterior mX neurons by delaying their axon formation. To test whether differences in RA signaling cause differences in the timing of axon initiation, we measured the timing of anterior and posterior axon initiation in embryos treated with either 50 nM RA, which induces high RA signaling throughout the nucleus (Fig. S1A-B), or with 1 µM of the RA-synthesis inhibitor DEAB, which reduces RA signaling throughout the hindbrain (Fig. S1C). Treatment was started at 24 hpf to avoid the early adverse effects of disrupted RA signaling on hindbrain patterning. At this stage vagus motor neurons are largely post-mitotic (Barsh et al., 2017). We expected that RA treatment would delay axogenesis in anterior mX neurons, and that DEAB would accelerate axogenesis in posterior mX neurons, and that consequently both treatments would decrease the time difference between anterior and posterior axogenesis. We assessed axogenesis timing by photoconverting posterior *Tg(isl1:Kaede)*-expressing mX neurons at 28 hpf and performing live imaging to compare when anterior (Kaede green) and posterior (Kaede red) axons appear. Surprisingly, we found that neither increasing nor decreasing RA levels had any effect on the difference in axogenesis timing between anterior and posterior neurons (Figs. 4A, S1D-F, Supplementary Movie 1).

The finding that RA does not control axon formation timing is intriguing, as it suggests that RA controls axon target selection through an as-yet-unidentified mechanism. To discover this mechanism, we again used RNA-seq, this time to identify candidate genes whose expression in mX neurons is regulated by RA. We treated *Tg(isl1:Kaede)* embryos with DMSO (control) or with 50nM RA starting at 24 hpf, dissected the posterior hindbrain at 38 hpf (when anterior, but not posterior, mX axons are innervating their targets), dissociated to single cell suspension, collected Kaede positive cells by flow cytometry, and sequenced 3 replicates of 4,500-8,300 cells per condition. Because axon targeting is mediated by chemoattractant and chemorepellent signals, we were particularly interested in identifying genes of this type in our RNA-seq dataset. Among the genes whose expression was significantly changed by RA treatment (397 up-regulated & 364 down-regulated genes), one of the most significant was *met*, which is down-regulated by RA treatment (3.1-fold decrease, FDR = 5.90E-10) (Fig. 4B). *met* encodes a receptor tyrosine kinase that is the only known receptor for Hgf (Bottaro et al., 1991). Hgf is a chemoattractant for spinal motor neurons (Ebens et al., 1996) and has been implicated in cranial motor neuron chemoattraction in hindbrain explant cultures (Caton et al., 2000), but a requirement for Hgf in topographic map development in general or in vagus development in particular has not been demonstrated. We validated our RNA-seq findings by performing *met* ISH in RA- and DEAB-treated embryos. As reported in the mouse (Kamitakahara et al., 2017; Wu and Levitt, 2013), zebrafish *met* is expressed in a subset of hindbrain motor neurons including the vagus motor nucleus. Importantly, we found that, at 38 hpf, *met* is normally restricted to the anterior part of the mX nucleus, and that the *met* expression domain is significantly reduced towards the anterior upon RA treatment and significantly expanded towards the posterior upon DEAB treatment (Fig. 4C-F). Thus, RA is an inhibitor of *met* expression within the mX nucleus. We additionally found that *hgfa* (one of two *hgf* genes in the zebrafish) is expressed in the pharyngeal arches, consistent with the possible role of Met-Hgf signaling for mX axon chemoattraction (Fig. 5A-C).

**Figure 4:**
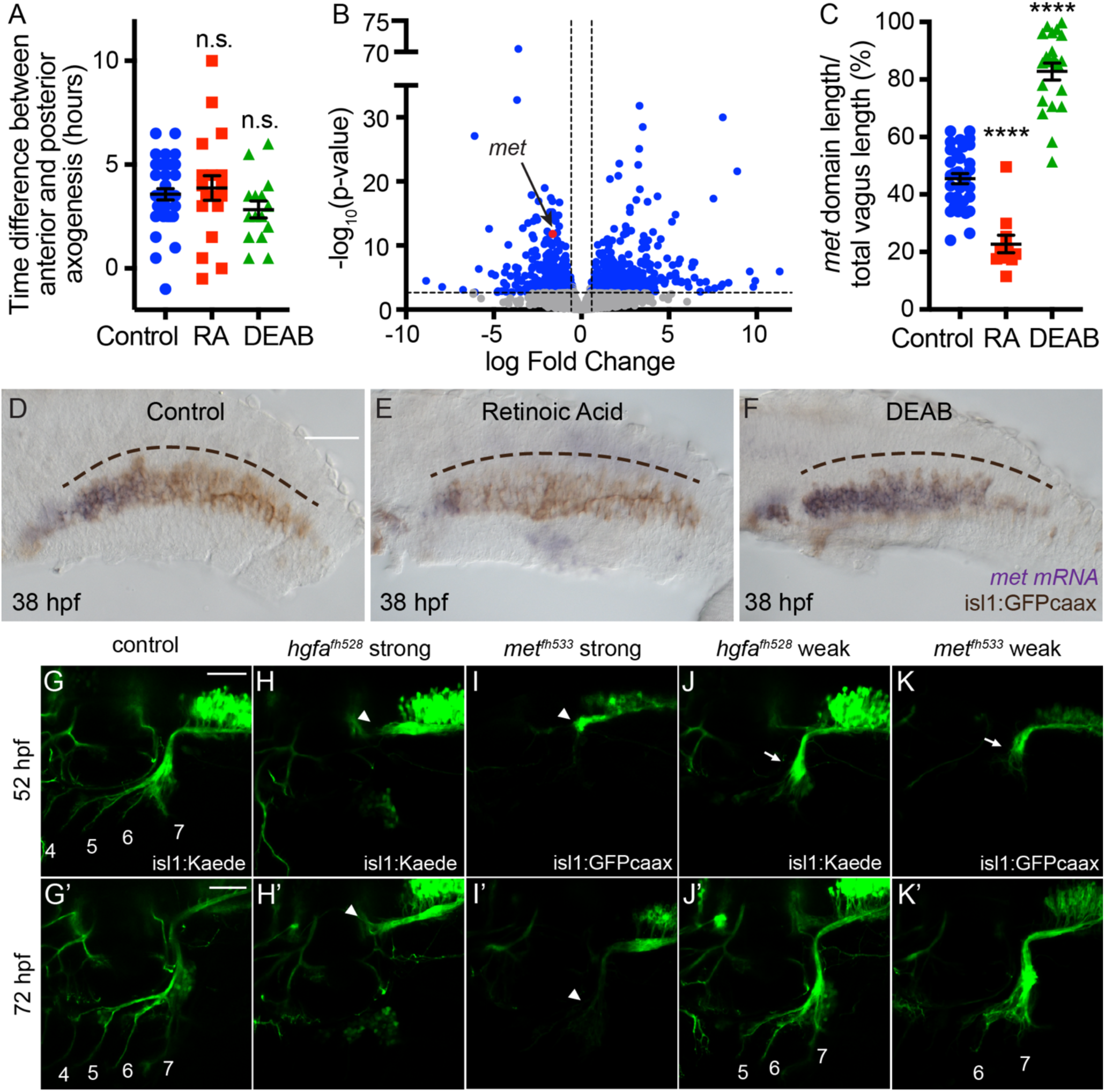
RA controls mX chemoattraction through Hgf/Met signaling, see also Supplementary Figures 1–2 & Supplementary Movies 1-3. (A) RA and DEAB treatment do not affect the difference in axogenesis timing between anterior and posterior mX neurons. Data represent mean ± SEM. t-test: ns = not significant. (B) Volcano plot of RNAseq data showing mRNAs depleted (left) or enriched (right) in RA-treated mX neurons relative to DMSO treated controls. Dashed lines indicate significance threshold for a false discovery rate of 5% (y-axis) and fold change of 1.5 (x-axis). Blue dots represent significantly differentially expressed genes. Red dot represents *met*. (C) Quantification of *met* expression domains at 38 hpf as represented in (D-F). Data represent mean ± SEM. t-test ****P<0.0001. (D-F) *met* RNA *in situ* hybridization in control (D), RA-treated (E) and DEAB-treated (F) embryos. mRNA expression is purple and the mX neurons marked by isl1:GFP are brown. The A-P span of the mX nucleus is indicated by the curved dotted line. (G-K) Paired 52hpf and 72hpf mX nerve structure in control (G), *hgfa^fh528^* (H, J), and *met^fh533^* (I,K) embryos. Phenotypes of varying strengths are seen. At 52hpf, in the strong phenotype mX axons (green) have stalled at the dorsal turning point (arrowhead) (H-I), while in the weaker phenotype axons have turned towards the PAs but have stalled at the ventral choice point (arrows) (J-K). In all cases, axons are deficient in entering the PAs at this stage whereas in controls axons have extended into PA4, 5 and 6 (G). At 72 hpf, in the strong phenotype axons remain stalled and fail to enter the PAs (H’-I’), while in the weak phenotype some axons escape and variably innervate PAs 5, 6, and 7 (J’-K’). Numbers indicate innervated PAs. All images are lateral views oriented with anterior to left. Scale bars = 50 μm.

**Figure 5:**
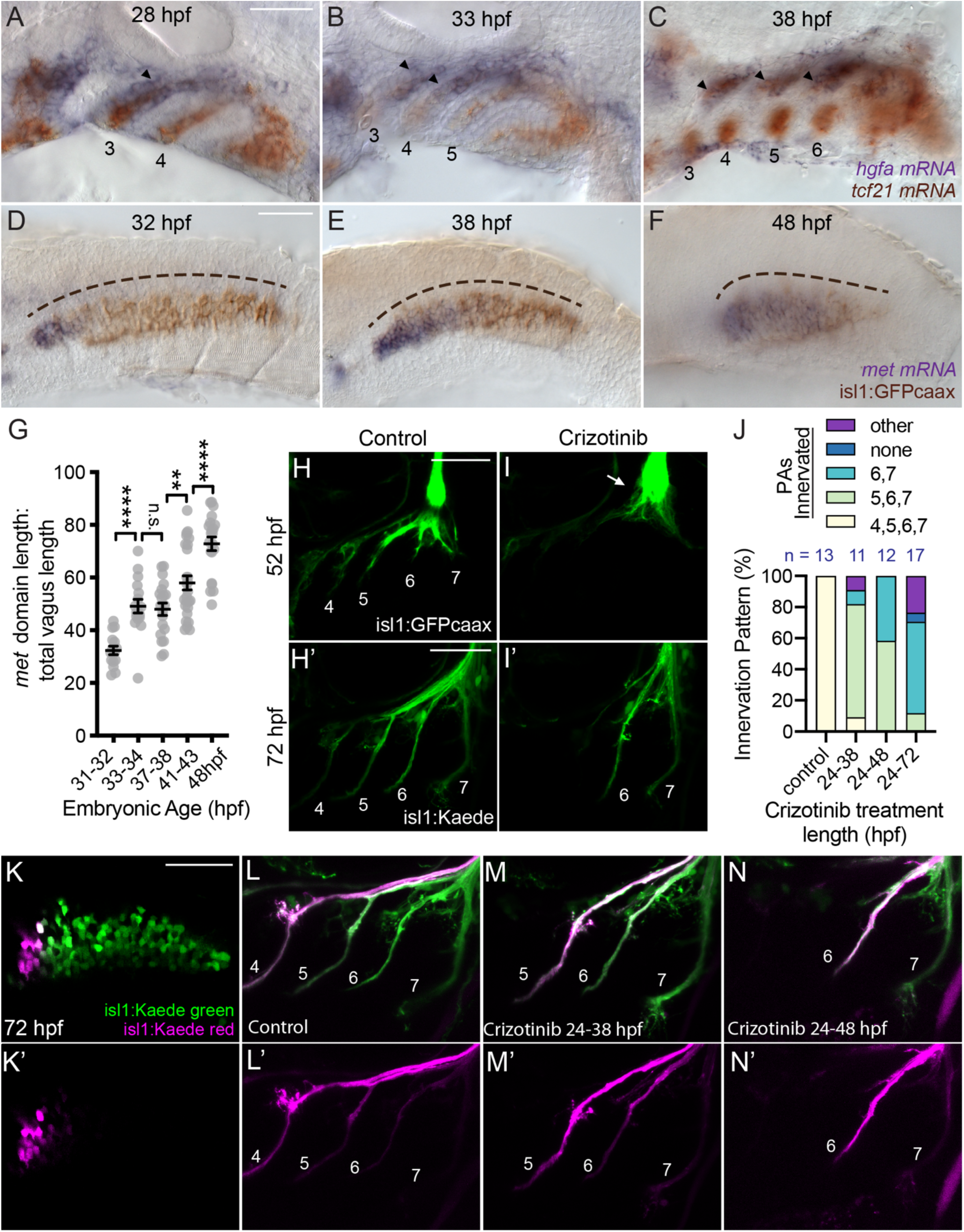
Spatiotemporal coordination of *hgfa* and *met* expression controls vagus motor axon target selection, see also Supplementary Figure 2. (A-C) Double *hgfa* (purple) and *tcf21* (brown) RNA *in situ* hybridization time series showing A-P sequential expression in the PAs. Arrowheads indicate *hgfa* expression in PAs. Numbers mark PAs. (D-F) *met* RNA *in situ* hybridization time series showing A-P expansion over time in the mX nucleus. mRNA expression is purple and the mX neurons marked by isl1:GFP are brown. The A-P span of the mX nucleus is indicated by the curved dotted line. (G) Quantification of *met* expression domains over time represented in (D-F). Data represent mean ± SEM. t-test ****P<0.0001, **P<0.005, ns = not significant. (H, I) mX nerve structure in DMSO-treated control (H) and Crizotinib-treated (I) embryos. At 52 hpf, mX axons (green) in Crizotinib-treated embryos stall at the dorsal choice point (arrow) (I). At 72 hpf, some mX axons escape and innervate posterior PAs (I’). Numbers indicate innervated PAs. (J) Quantification of arch innervation pattern frequency represented in (H-I & K-N). Data represent the frequency with which axons (red or green) are observed in each branch. Progressively longer Crizotinib treatments cause progressively more severe loss of anterior branches. (K-N) Delayed Hgf/Met signaling causes a timing-dependent posterior shift in anterior mX axon target selection. Numbers indicate PAs. (K) Representative anterior mX neuron photoconversion used for anterograde labeling in (L-N). (L) mX branches in control DMSO-treated control embryo. Anterior mX neurons predominantly target PA4. (M) mX branches in embryo treated with Crizotib from 24-38 hpf. The PA4 branch is missing, and anterior mX neurons predominantly target PA5. (N) mX branches in embryo treated with Crizotib from 24-48 hpf. The PA4 and PA5 branches are missing, and anterior mX neurons predominantly target PA6. All images are lateral views oriented with anterior to left. Scale bars = 50 μm.

Do Met and Hgf mediate mX axon chemoattraction? To test this, we generated frameshift alleles of *met* (*met^fh533^)* and *hgfa* (*hgfa^fh528^* and *hgfa^fh529^*) and examined their effect on mX axon targeting. We also examined the previously described *met^s908^* hypomorphic allele (Anderson et al., 2013). In these mutants, although PA formation and mX neuron differentiation occur normally (Fig. S2A-B, Supplementary Movies 2 & 3), PA innervation is strongly disrupted. The phenotypes we observed vary in severity; the strongest phenotype, observed in all alleles except *met^s908^*, is a complete loss of all vagus motor innervation of the PAs and viscera. In these cases, mX axons extend anteriorly towards the otic vesicle but fail to turn ventrally towards the PAs (Figs. 4G-I, S2D-E). All alleles also exhibited a weaker innervation defect in some embryos, in which axons exhibit a prolonged stall at the branch point (often until 56hpf or later) but eventually attain partial innervation (Figs. 4J-K, S2F-G). In these cases, branch formation is incomplete – the PA4 branch never forms and the PA 5, 6, and 7 branches sometimes fail to form – and the branches that do form do so late and often appear thinner than normal. These findings reveal that Hgf, signaling through Met, is the chemoattractant that guides mX axons to their PA targets.

### Spatiotemporal coordination of *hgfa* and *met* expression controls vagus motor axon target selection

Although *met* is necessary for mX axons to innervate the PAs normally, it is not clear whether or how it might instruct the differential targeting decisions that underlie topographic organization. One potential explanation is that spatially graded *met* receptor expression promotes differential target selection. However, we have previously observed that the timing of axon targeting can override spatial identity in determining axon target selection (Barsh et al., 2017). Therefore, we considered that topographic mX axon target selection may result from differences in the timing, rather than the level, of chemoattractant signaling along the A-P axis. To examine whether the spatial pattern of Hgf/Met signaling changes over time, we observed the expression of these genes by ISH over the course of axon targeting. *hgfa* mRNA appears in each arch shortly after that arch appears, based on the emergence of a distinct tcf21:mCherry muscle precursor core (Fig. 5A-C), resulting in an anterior to posterior sequence of *hgfa* expression that reflects the sequential formation and subsequent innervation of the PAs (Fig. 1C-D) (Barsh et al., 2017). Intriguingly, we saw a corresponding progression of *met* expression in the mX nucleus. At 32 hpf, the stage when the axons of anterior-most mX neurons are entering PA4 (Fig. 1D), *met* mRNA is restricted to the anterior-most region of the mX nucleus (Fig. 5D). *met* expression then steadily expands posteriorly to fill the majority of the nucleus by 48hpf (Fig. 5D-G). The expansion of the *met* expression domain corresponds with a recession of the inhibitory RA signaling domain, which can be inferred from *hoxb5a* mRNA expression. *Hoxb5a* is directly regulated by RA (Emoto et al., 2005; Oosterveen et al., 2003) and we observe that its anterior limit in the vagus region recedes between 24 hpf and 32 hpf (Fig. S3A-D). The 6-hour delay in *met* expansion compared to *hoxb5a* recession likely reflects delays in the degradation of an as-yet unidentified RA-dependent *met* repressor. We were unable to detect RA recession with the 12xRARE:GFP reporter, which is less sensitive to RA than the endogenous *hoxb5a* gene and thus is never expressed in the anterior vagus region.

The above findings suggest that the domain of Hgf/Met signaling in the vagus motor nucleus is temporally constrained, and suggest a model for how topographic targeting is controlled, which we refer to as “temporal matching”. The correlation between the sequential onset of *hgfa* expression in the PAs and their sequential innervation suggests that *hgfa* expression dynamics function to establish distinct developmental windows during which each PA is able to attract mX axons. Likewise, the expanding *met* expression domain in the mX nucleus functions to establish a distinct developmental time at which each neuron becomes competent to respond to the Hgf signal. Thus, the earliest *met*-expressing mX neurons (the anterior-most) become temporally matched with the earliest *hgfa*-expressing PA (PA4). As *met* and *hgfa* expression domains extend posteriorly, sequentially more posterior neurons become matched to sequentially more posterior PAs. We propose that the RA signal determines axon target selection by controlling the pace of expansion of the *met* expression domain and, therefore, which mX regions become temporally matched to each PA.

The temporal matching model makes a key prediction: that delaying Hgf/Met signaling in anterior mX neurons will be sufficient to shift targeting of their axons to more posterior branches, and that the extent of the posterior shift will be proportional to the length of the delay of signaling. To test this hypothesis, we delayed Hgf/Met signaling with Crizotinib, a small molecule that inhibits Met tyrosine kinase activity by specific and reversible competitive binding to the ATP-binding pocket (Zou et al., 2007). Treatment of embryos with Crizotinib starting at 24hpf consistently recapitulates the less severe *met^fh533^* and *hgfa^fh528^* mutant phenotypes, resulting in a prolonged stall of mX axons at the branch point (Fig. 5H-I). However, unlike in the mutants, Met function can be rapidly restored by washing out the drug, thus allowing us to experimentally control the onset of signaling. We treated embryos with Crizotinib starting at 24hpf and washed out the drug at two timepoints – 38 hpf, corresponding to the time that PA5 innervation normally begins, and 50 hpf, corresponding to the time that PA6 innervation normally begins. We then allowed axon targeting to proceed until 72hpf and assessed the targeting of anterior mX neurons by Kaede photoconversion. We observed that delaying Hgf/Met signaling causes progressive loss of anterior PA branches. Delaying signaling until 38 hpf prevents the PA4 branch from forming, while delaying signaling until 50 hpf additionally frequently prevents the PA5 branch from forming (Fig. 5J-N); these phenotypes are weaker than when we do not wash out the drug (Fig. 5J).

The loss of anterior mX branches after Crizotinib treatment could be because anterior mX neurons are prevented from innervating any PAs, or because they are redirected to more posterior PAs, the latter result being more consistent with the temporal matching hypothesis. Using Kaede photoconversion of the anterior-most 10% of mX neurons, which normally innervate PA4 and PA5, we found that these neurons are indeed redirected to the next most anterior PA that forms:PA5 in the 38 hpf washout or PA6 in the 50 hpf washout (Fig. 5K-N). These results show that controlling the timing of Hgf/Met signaling is sufficient to determine mX axon target selection, and confirm our model that mX axon target selection is driven by coordinating the spatiotemporal dynamics of *met* expression in the mX nucleus with the sequential differentiation of the PAs, thus establishing the topographic innervation pattern of the vagus nerve (Fig. 6).

**Figure 6:**
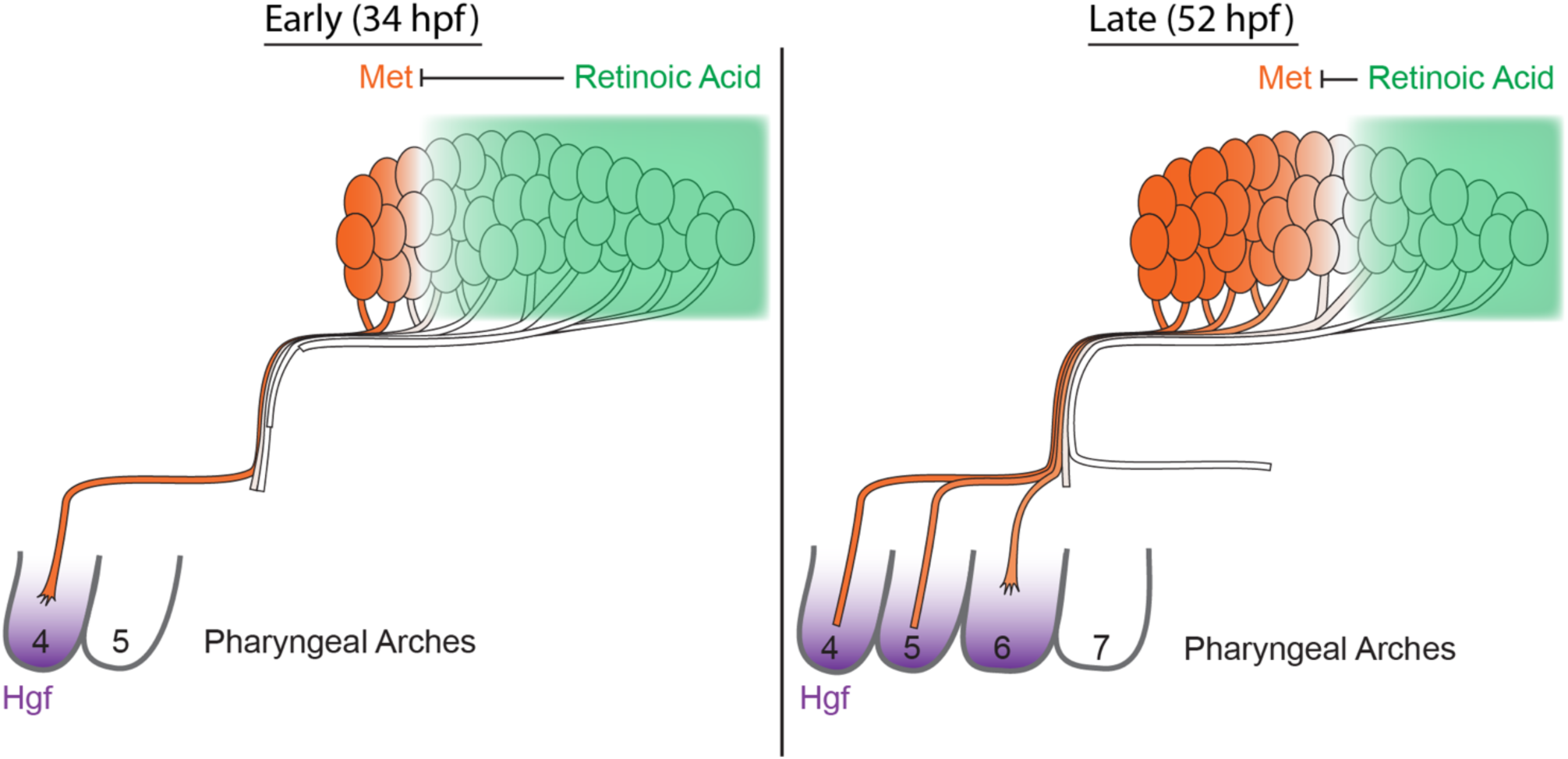
Temporal matching model of topographic vagus motor axon targeting. During early innervation (left), the anterior-most PA (PA4) forms and turns on Hgf, becoming competent to accept mX axons. At the same time, a broad RA signaling domain in the mX nucleus represses *met* expression in all but the most anterior mX neurons, which are thus competent to respond to PA4-derived Hgf chemoattraction. Later (right), more posterior PAs have formed and turned on Hgf, thus becoming competent to accept mX axons. At the same time, recession of the RA signaling domain in the mX nucleus derepresses *met* expression in more posterior mX neurons, making them competent to respond to more posterior PA-derived Hgf signal.

## DISCUSSION

Topographic patterning is important for the coherent flow of information through complex neural circuits, but presents a considerable organizational challenge in the developing nervous system. Here, we elucidate how precise axon targeting decisions are regulated in the zebrafish embryo to promote topographic organization of the branched vagus motor nerve. We first quantified the spatiotemporal dynamics of mX branch formation, which occurs in an anterior-to-posterior sequence. We then found that differential RA signaling in the mX nucleus specifies positional identity and showed that mX axon target selection is autonomously determined by the level of RA signaling in mX neurons. Next, we discovered that RA regulates *met* receptor expression dynamics in the mX nucleus and that Met and its PA-derived ligand, Hgf, constitute the chemoattractant signal that guides mX axons to the PAs. Finally, we showed that manipulating the timing of Hgf/Met signaling is sufficient to alter mX axon target selection. We present a new temporal matching model of topographic map development in which an RA-dependent wave of Hgf/Met signaling moving across the A-P axis of the vagus motor nucleus drives topographic axon target selection (Fig. 6).

The classic chemoaffinity model of topographic axon targeting posits that the communication of spatial identity differences between neurons and their targets guides axon targeting (Cang and Feldheim, 2013; Sperry, 1963). While this model adequately explains topographic map development in some systems, such as the retinotopic map, in some cases spatial signaling alone is insufficient to direct the requisite targeting decisions (Eerdunfu et al., 2017; Kulkarni et al., 2016; Petrovic and Hummel, 2008; Takeuchi et al., 2010). Here, we present a new model of topographic targeting, which we refer to as temporal matching, in which chemoattractant signaling between neurons and their targets is coordinated in both space and time to precisely guide axon targeting (Fig. 6). In the target region, the PAs form and express *hgfa* sequentially from anterior to posterior, causing each PA to become accessible to mX innervation at a different time. Meanwhile, in the mX nucleus, an anterior to posterior wave of *met* expression causes neurons to become competent to respond to Hgf at different times. The result of these two independent events is that, at any given point in time, neurons at a particular A-P position are guided to a particular PA. The temporal matching model represents a new mechanism of topographic map development by which neurons can integrate both spatial and temporal cues to make complex axon targeting decisions.

We identify a critical role for RA signaling in mX neuron identity. RA is a well-established regulator of early rhombomeric A-P patterning in the hindbrain (Moens and Prince, 2002; Nolte et al., 2019) and, later, in the specification of motor neuron subtypes in the spinal cord (Sockanathan and Jessell, 1998; Sockanathan et al., 2003). RA signaling has also been proposed to provide positional information for topographic axon targeting in the olfactory and retinotopic maps (Login et al., 2015; Sen et al., 2005). In these cases, however, RA is believed to work in a spatially graded manner to establish differential cell identities at different levels of RA signaling. Its role in the mX nucleus is fundamentally different in that there is no indication that RA acts in a spatially graded manner; rather, it appears that differences in the *time* at which neurons are relieved from RA signaling is what determines topographic mapping. These observations indicate the risks of inferring signal function from a single snapshot in time, and the importance of considering temporal, as well as spatial, dynamics when studying RA, and other signaling molecules, during embryonic patterning.

This work also represents an important advance in our understanding of the developmental role of Hgf/Met signaling. Hgf/Met was one of the first motor axon chemoattractants to be discovered, and has long been known to promote motor neuron outgrowth *in vitro* and *in vivo* (Caton et al., 2000; Ebens et al., 1996). Since this discovery, the detailed developmental mechanisms of several other major axon guidance signals has been elucidated (Bellon and Mann, 2018; Robichaux and Cowan, 2014; Stoeckli, 2017), while our understanding of how Hgf/Met regulates axon targeting decisions in the embryo has advanced comparatively little. This may be in part because in the limb, where the role of Hgf/Met signaling has been best studied, it has multiple successive functions: in the migration of muscle precursors into the limb (Bladt et al., 1995; Brand-Saberi et al., 1996; Haines et al., 2004; Nord et al., 2019; Talbot et al., 2019), in the formation of specific limb-innervating motor neuron pools (Helmbacher et al., 2003), in motor axon attraction (Ebens et al., 1996), and in motor neuron survival (Lamballe et al., 2011; Novak et al., 2000; Yamamoto et al., 1997). Hgf/Met is not required for the specification or survival of mX neurons, and although recent work has shown that Hgf/Met signaling is required for the migration of posterior PA-derived muscle progenitors to form esophageal muscle in the mouse (Comai et al., 2019), we do not observe any abnormalities in the formation of posterior PA muscle progenitors in the zebrafish *hgfa* or *met* mutants at the stages we studied here. Therefore, we have been able to study its function specifically in motor axon guidance. We find that precise developmental regulation of signaling between Hgf and Met plays an essential and direct role in guiding the topographic targeting of vagus motor axons. Whereas in the limb, specificity of Hgf/Met dependent innervation is determined by *which* motor neuron pools express Met, the specificity of the vagus topographic map depends on *when* vagus motor neurons express Met.

A key aspect of topographic mX nerve formation is that posterior mX neurons are delayed in the innervation of their targets. In this paper, we propose that the delayed onset of *met* expression causes this delay and show that experimentally delaying Hgf/Met signaling is sufficient to shift axon target selection to more posterior PAs. We have also previously reported that posterior mX neurons form axons several hours later than anterior mX neurons (Barsh et al., 2017). This phenomenon could also cause the requisite delay in axon target selection and, in support of this idea, we have shown that delaying axon formation is sufficient to direct axons to more posterior PAs. Therefore, two events – onset of *met* expression and initiation of axon formation – are independently regulated to control mX axon targeting. Why do two such apparently redundant mechanisms exist? It may be that each is required, but not sufficient, to generate the full length of delay required for posterior targeting. Indeed, the difference in timing between PA4 and PA7 innervation (20 hours) is longer than either the difference in timing of axogenesis or the difference in timing of met expression by PA4 and PA7-targeting axons. Furthermore, neither delaying axogenesis nor delaying Hgf/Met signaling is sufficient to completely disrupt topographic mapping (Barsh et al., 2017). It is therefore likely that these two mechanisms cooperatively promote temporal matching in the vagus nerve.

Our model raises several questions for future investigation. 1) How does RA signaling repress *met* expression? While RA is generally believed to activate the transcriptional activity of its nuclear receptors (Evans and Mangelsdorf, 2014; Ghyselinck and Duester, 2019), whether it represses *met* directly, or indirectly via induction of a repressor, remains to be seen. *hox5* genes are directly RA-regulated (Oosterveen et al., 2003), and we previously described a role for *hox5* genes in promoting the targeting of mX axons to posterior PAs (Barsh et al., 2017). However, *met* expression is unaffected in *hoxa5a; hoxb5a; hoxb5b* triple mutants that we have generated (unpublished data). Thus, while RA regulates both the Hgf/Met-dependent and *hox5*-dependent aspects of mX guidance, the two mechanisms are not interdependent. We anticipate that the *met* repressor will be found amongst the RA-upregulated transcripts in our vagus RNA-Seq dataset. 2) What sets the PA innervation windows? The onset of *hgfa* expression appears to initiate PA innervation, but what determines when axons switch to target the next PA? In other contexts, competition between axons can cause shifts in target selection over time, for instance by the innervation-induced expression of a chemorepellant (Takeuchi et al., 2010); however, our observation that delaying Hgf/Met signaling causes anterior axons to switch to more posterior targets without prior innervation of more anterior PAs indicates that switching between PA targets is independent of innervation. While *hgfa* mRNA is not downregulated in the PAs over time, downregulation of a required processing factor (Kawaguchi and Kataoka, 2014) or the delayed expression of a repulsive or inhibitory cue could cause axons to switch targets. Alternatively, the ongoing morphogenesis that moves the PAs anteriorly relative to the brain, which can be seen in time-lapse movies (Supplementary Movies 2 & 3), could make posterior PAs more easily accessible over time. The innervation-independent switching mechanism is a key component of topographic mapping that remains to be discovered. 3) How is developmental timing coordinated between the mX nucleus and the PAs? Our experiments show that misaligning the timing of PA formation and mX axon targeting has severe consequence on the topographic map, but how is the alignment of these distant tissues normally maintained? Is timing in each tissue independently controlled, or does a common signal regulate the development of both tissues? RA is essential for multiple aspects of PA development (Kopinke et al., 2006; Li et al., 2012; Linville et al., 2009; Mark et al., 2004; McGurk et al., 2017) and changes in RA signaling are thought to contribute to the craniofacial defects in DiGeorge syndrome (Karpinski et al., 2014). Whether RA controls the timing of PA emergence and/or *hgfa* expression, thereby entraining PA development to RA-regulated events in the vagus motor nucleus, remains to be determined.

A major challenge in developmental biology is how to precisely generate varied and complex structures in a small space with a limited molecular toolkit. This challenge is particularly acute in the nervous system, in which the formation of intricate innervation patterns requires the communication of diverse spatial and identity information through the noisy embryonic environment. Many innovations in developmental complexity have been built on increases in signal complexity – the ability to interpret more varied and combinatorial signals along more embryonic axes with higher spatial resolution. It is in this context that most topographic mapping events have been interpreted. Our work reveals that the communication of information along the temporal axis is also crucial in generating the signal complexity required for topographic axon targeting, thus highlighting the importance of developmental time as an additional dimension with which to increase signal resolution. The importance of combining spatial and temporal information in generating the complexity of the nervous system is evident in other contexts as well. For instance, temporal transcription factor cascades in neural progenitors allow for the generation of a diversity of cell types greater than could be achieved by spatial cues alone (Doe, 2017; Holguera and Desplan, 2018). Likewise, differences in the timing of neuron birth can direct differential axon targeting decisions (Eerdunfu et al., 2017; Petrovic and Hummel, 2008; Takeuchi et al., 2010). However, to the best of our knowledge, the vagus nerve represents the first example of a temporal axon targeting mechanism that is independent of cell birth timing (Barsh et al., 2017). While temporal signaling dynamics are inherently more difficult to identify and study than spatial signaling dynamics, it is likely that the temporal axis is a major, and underrepresented, contributor to the organization of the nervous system, and the embryo as a whole.

## ACKNOWLEDGEMENTS

We thank Rachel Garcia for excellent animal care. Victoria Prince, Ken Poss, Josh Waxman, Jeremy Rabinowitz, Ryan Anderson, and Chuck Kimmel generously provided transgenic lines, constructs, and reagents. We thank Jeff Delrow and Ryan Basom for assistance with RNAseq, and Tim Randolph for statistical assistance. We thank all members of the Moens lab for discussion and editing. Funding for this project was provided by NIH grant R01 NS109425 to C.B.M., F32 HD096860 to A.J.I., and F30 NS093703 to G.R.B., as well as American Heart Association grant 18POST33990492 to A.J.I.

## AUTHOR CONTRIBUTIONS

Conceptualization, A.J.I., G.R.B., C.B.M.; Methodology, A.J.I., G.R.B., C.B.M.; Software, A.J.I.; Formal Analysis, A.J.I.; Investigation, A.J.I., G.R.B., J.A.S, C.B.M.; Writing – Original Draft, A.J.I., C.B.M.; Writing – Review and Editing, A.J.I., G.R.B., J.A.S., C.B.M.; Visualization, A.J.I., J.A.S., C.B.M.; Supervision, A.J.I., C.B.M.; Project Administration, A.J.I., C.B.M.; Funding Acquisition, A.J.I., G.R.B., C.B.M..

## STAR METHODS

### Lead Contact and Materials Availability

Further information and requests for resources and reagents should be directed to and will be fulfilled by the Lead Contact, Cecilia Moens.

### Experimental model and subject details

#### Zebrafish care and maintenance

*Danio rerio* animals were raised at the Fred Hutchinson Cancer Research Center facility in accordance with IACUC-approved protocols. All experiments were carried out in accordance with IACUC standards. Fish were bred and maintained according to standard protocols (Westerfield, 2000). All experimental stages are noted in the figures and text. Staging of embryos between 24-38 hpf was done according to the prim staging method (Kimmel et al., 1995). Staging of embryos beyond 38 hpf was done by elapsed time at 28°C. Sex is not a relevant biological variable in our experiments, as they are carried out before sex is determined in zebrafish (Siegfried, 2010). Transgenic lines used in this study include *Tg(isl1-hsp70l:Kaede)* (Barsh et al., 2017); *Tg(isl1:Gal4)fh452* (Davey et al., 2016); *Tg(isl1:mRFP)fh1* (Grant and Moens, 2010); *TgBAC(tcf21:mCherry-NTR)pd108* (Wang et al., 2015); *Tg(isl1:eGFPCAAX)fh474* (Barsh et al., 2017); *Tg(12xRAREef1a:GFP)sk71* (Waxman and Yelon, 2011). Mutant lines used in this study include *met^s908^* (Anderson et al., 2013) and *cyp26a1^rw716^* (Emoto et al., 2005). Mutant lines generated for this study include *met^fh533^*, *hgfa^fh528^*, and *hgfa^fh529^* (see details below).

### Method Details

#### Plasmid construction and injection

The following plasmids were generated for this study: 10X-UAS:eGFP-CAAX-polyA, 10X-UAS:DN-hRARa-p2a-eGFP-CAAX-polyA, and 10X-UAS:CA-RARga-p2a-eGFP-CAAX-polyA. The 10X-UAS and eGFP-CAAX sequence were obtained from the Tol2kit (Kwan et al., 2007). Plasmids containing the DN-hRARα and CA-RARga were kindly provided by the Waxman lab (Waxman and Yelon, 2011) and cloned using the following primers:

> DN-hRARα: Forward ATGGCCAGCAACAGCAGCTC
>
> Reverse CGGGATCTCCATCTTCAGCG
>
> CA-RARga: Forward ATGGCCCCCCCGACCGATG
>
> Reverse CTGAGCTCTTCCTCCGTGGC

RAR sequences were inserted into a pDONR 221 vector in-frame upstream of a p2a-eGFP-CAAX element (Barsh et al., 2017). Final constructs were assembled in a pBHR4R3 plasmid (gift of the Brockerhoff lab) using the Gateway system (Life Technologies). Sparse transgenic labeling of cells was done by injecting one-cell-stage embryos with 50pg of plasmid and 50pg of Tol2 mRNA, and screening embryos for sparse GFP expression at 2 days post fertilization on a Zeiss AxioZoom.V16 microscope.

#### Generation of *met* and *hgfa* mutant alleles

*met* and *hgfa* mutant alleles were generated using CRISPR/Cas9 as described in (Talbot and Amacher, 2014) using the short oligo method with the following modifications: gRNAs were designed using chopchop (https://chopchop.cbu.uib.no). One-cell-stage embryos were injected with 200pg gRNA and 500pg Cas9 protein, and F1 mutant animals identified by sequencing. The following gRNA targeting sequences were synthesized and co-injected:

> *met* gRNA 1 GGTTCTGGCCATCTGGCTCG
>
> gRNA 2 GGCTTCGGCTGCGTGTTTCA
>
> *hgfa* gRNA 1 GGAGTGTATGAAATGTAATG
>
> gRNA 2 TTCGGCAAAAGTTCTGACGT
>
> gRNA 3 TCGTCCGTGGTGTTACACGA

*met^fh533^* is a 7bp deletion starting at nucleotide 3,280, and a 17bp insertion at nucleotide 3,301 of the *met* coding sequence, resulting in a frameshift at amino acid 1094 and a premature stop at amino acid 1099. *hgfa^fh528^* is a 79bp deletion covering the start of exon 6. *hgfa^fh529^* is a 11bp insertion at nucleotide 474, and a 265 bp deletion at nucleotide 597 of the *hgfa* coding sequence, resulting in a frameshift at amino acid 159 and a premature stop at amino acid 201.

#### RNA *in situ* hybridization

Anesthetized embryos were fixed overnight at 4°C in 4% paraformaldehyde with 1X PBS and 4% sucrose. RNA *in situ* hybridization was performed as described in (Prince et al., 1998). GFPcaax (from *Tg(isl1:GFPcaax)*) protein was visualized in tandem with mRNA signal as follows. After completion of *in situ* hybridization, embryos were washed and incubated with primary antibody (rabbit anti-GFP). Samples were re-blocked for 1 hour, stained with secondary antibody (mouse anti-rabbit HRP, 1:200), washed 3x with PBS+0.1% Tween 20 and color was developed with DAB solution. For imaging, embryos were cleared step-wise into 75% glycerol, dissected, and imaged on a Zeiss Axioplan2 microscope. The *hoxb5a* probe is from (Prince et al., 1998). The *met* and *hgfa* probes are from (Haines et al., 2004). The tcf21 probe is from Chuck Kimmel and can be amplified with the following primers:

> *tcf21*: Forward CGTTTCCACATAGCCAGTTGC
>
> Reverse GGAGAGTTTGGTGTCCGGCGG

The *crabp1b* and *dhrs3b* probes were amplified from cDNA prepared from 2dpf embryos using the following primers and *in vitro* transcribed with the MEGAscript T7 transcription kit.

> *crabp1b*: Forward TGCTGAGAAAAGTGGCTTGTGC
>
> Reverse taatacgactcactatagggagaCACATCATCAGCCCCAAACATC
>
> *dhrs3b*: Forward CGGACGGAAAAATGTCTGAAGG
>
> Reverse taatacgactcactatagggagaGATAACGAGTGCGTTCATGGTCC

#### Fluorescence microscopy

All embryos were anesthetized with 400mg/L MESAB, embedded in 1.4% (for single time point imaging) or 0.7% (for time-lapse imaging) low-melt agarose and imaged live on a Zeiss LSM 700 confocal microscope at the stages noted. Images were processed using imageJ (Schneider et al., 2012). For Kaede photoconversion, the appropriate region was photoconverted using 200 iterations of the 405 laser at 10% power.

#### RNA sequencing

To prepare embryos for sequencing of anterior and posterior mX neuron populations (A-P experiment), 28 hpf *Tg(isl1:Kaede)* embryos were mounted for fluorescence microscopy and cells in the anterior or posterior 25-30% of the mX nucleus were photoconverted as described above, and unmounted. To prepare embryos for sequencing of DMSO- and RA-treated mX neuron populations (RA experiment), *Tg(isl1:Kaede)* embryos were treated with drugs as described below and dissected at 38 hpf. For both experiments, hindbrain rhombomere 8 was dissected in calcium-free Ringer’s solution with MESAB, and dissected tissue was dissociated to single-cell suspension by pipetting in 0.25%Trypsin-EDTA for 5 minutes. For the A-P experiment, dissociated cells transferred into cold DPBS + 5% FBS + 1% BSA. The cell suspension was then transferred onto a glass coverslip, covered with mineral oil, and photoconverted cells were manually picked up using a 10μm diameter transplant pipette on a hydraulic micromanipulator mounted on a Zeiss AxioSkip microscope. Picked cells were transferred into lysis buffer from the RNA isolation kit (see below) and frozen at −80°C until RNA isolation. For the RA experiment, dissociated cells were transferred into cold DPBS + 1%BSA + 2μg/mL DAPI. Cells were then sorted on an BD FACS ARIA II flow cytometer. Kaede+, DAPI-cells were collected in lysis buffer from the RNA isolation kit and immediately processed. For both experiments, RNA was isolated using the RNAqueous-Micro Total RNA Isolation Kit, cDNA was amplified using the SMART-Seq v4 Ultra Low Input RNA Kit for Sequencing, and libraries were prepared using the Nextera XT DNA library prep kit and sequenced on an Illumina HiSeq 2500 sequencer. For the A-P experiment, 3 replicates of 100 cells were sequenced and analyzed for each position. Reads of low quality were filtered prior to alignment to zebrafish (GRCz10) using TopHat v2.1.0 (Trapnell et al., 2009). Counts were generated from TopHat alignments for each gene using HTSeq v0.6.1, (Anders et al., 2015), employing the “intersection-strict” overlap mode. Genes with low counts across all samples were removed, prior to identification of differentially expressed genes using the Bioconductor package edgeR v3.18.1 (Robinson et al., 2009). A false discovery rate (FDR) method was employed to correct for multiple testing (Reiner et al., 2003), with differential expression defined as |log2 (ratio) | >= 0.585 (± 1.5-fold) with the FDR set to 10%. For the RA experiment, 3 replicates of 4500-10000 cells were sequenced and analyzed for each condition. Reads of low quality were filtered prior to alignment to zebrafish (GRCz11) using STAR v2.5.2a (Dobin et al., 2013) in 2-pass mode. Counts were generated from were generated from STAR alignments for each gene using featureCounts from the Subread package v1.5.0 (Liao et al., 2014). Genes with low counts across all samples were removed, prior to identification of differentially expressed genes using the Bioconductor package edgeR v3.20.9 (Robinson et al., 2009). A FDR method was employed to correct for multiple testing (Reiner et al., 2003), with differential expression defined as |log2 (ratio) | >= 0.585 (± 1.5-fold) with the FDR set to 5%.

#### Drug treatments

Embryos were routinely treated with 0.2mM PTU at 24hpf to block pigment formation. For RA and DEAB treatments, embryos were dechorionated and treated with 50nM RA in DMSO or 1μM DEAB in DMSO, or the equivalent volume of DMSO as a control, beginning at 24hpf. For Crizotinib treatments, embryos were dechorionated and treated with 20μM Crizotinib in DMSO, or the equivalent volume of DMSO as a control, beginning at 24hpf. For Crizotinib wash-out, embryos were transferred to fresh water at the specified times.

### Quantification and Statistical Analysis

Unless otherwise noted, all image analysis was performed in imageJ, and all statistics were performed in GraphPad Prism in the form of unpaired t-tests. Additional statistical information on each experiment is noted in figure legends.

#### Quantification of Axon Initiation Timing

We started at the end of the movie when axons were clearly visible, and tracked backwards until the first frame in which the axon was visible. The green (un-photoconverted) channel was used to assess anterior mX neurons, and the red (photoconverted) channel was used to assess posterior mX neurons.

#### Quantification of branch distribution of axons from anterior mX neurons

At 72hpf, neurons in the anterior 10% of the mX nucleus were photoconverted and 1 hour allowed for diffusion of converted protein into axons before imaging of axon branches. In ImageJ, from maximum intensity projection images of axon branches, a 3.9μm-thick line was drawn that covered all four PA branches (see Fig. 3A) and Plot Profile tool was used to measure green and red intensity values along that line. A custom Python script was then used to identify the intensity peaks in the green channel that correspond to the axon branches, and to measure the sum of the red and green fluorescence values under those peaks. From these values, the normalized red fluorescence value for each peak was calculated as (red/(red+green)).

#### Quantification of mX nerve topography using sparse GFP labeling

F0 transgenic embryos were generated as described above, sorted for very sparse GFP labeling of mX neurons, and imaged as in Fig. 3C. To measure the A-P position of the labeled cell, the mX nucleus was manually divided into 10 bins across the A-P axis and the bin in which that cell was contained was noted. The branch which that axon innervated was also noted. In some cases, more than one cell was labeled by GFP; in these cases, we measured the position of the anterior-most GFP+ neuron, and the position of the anterior-most GFP+ axon. This data is graphed as position probability matrices in which the height of numbers on the Y-axis represents the percentage of cells in that position that innervate the indicated branch. We tested whether the target distribution at each position in DN-RAR and CA-RAR was significantly different from that of control using a Fisher’s exact test in R. Posterior regions 5-10 were pooled for this analysis to increase power because we observed low numbers of cells in these experiments.

#### Quantification of mRNA expression domain size

In RNA *in situ* hybridization images, we measured the length of a line drawn from the anterior edge of the mX nucleus to the posterior boundary of mRNA expression, and divided this by the measured length of the mX nucleus.

### Data and Code Availability

Genomic data generated in this study has been deposited to the NCBI Gene Expression Omnibus (GEO) with the following accession numbers: GSE135780 for the A-P experiment, and GSE135781 for the RA experiment. Custom code used in this study has been deposited in the public GitHub repository MoensLab/Isabella_et_al_2019.

## SUPPLEMENTAL INFORMATION

**Supplementary Figure 1, related to Figure 4.**
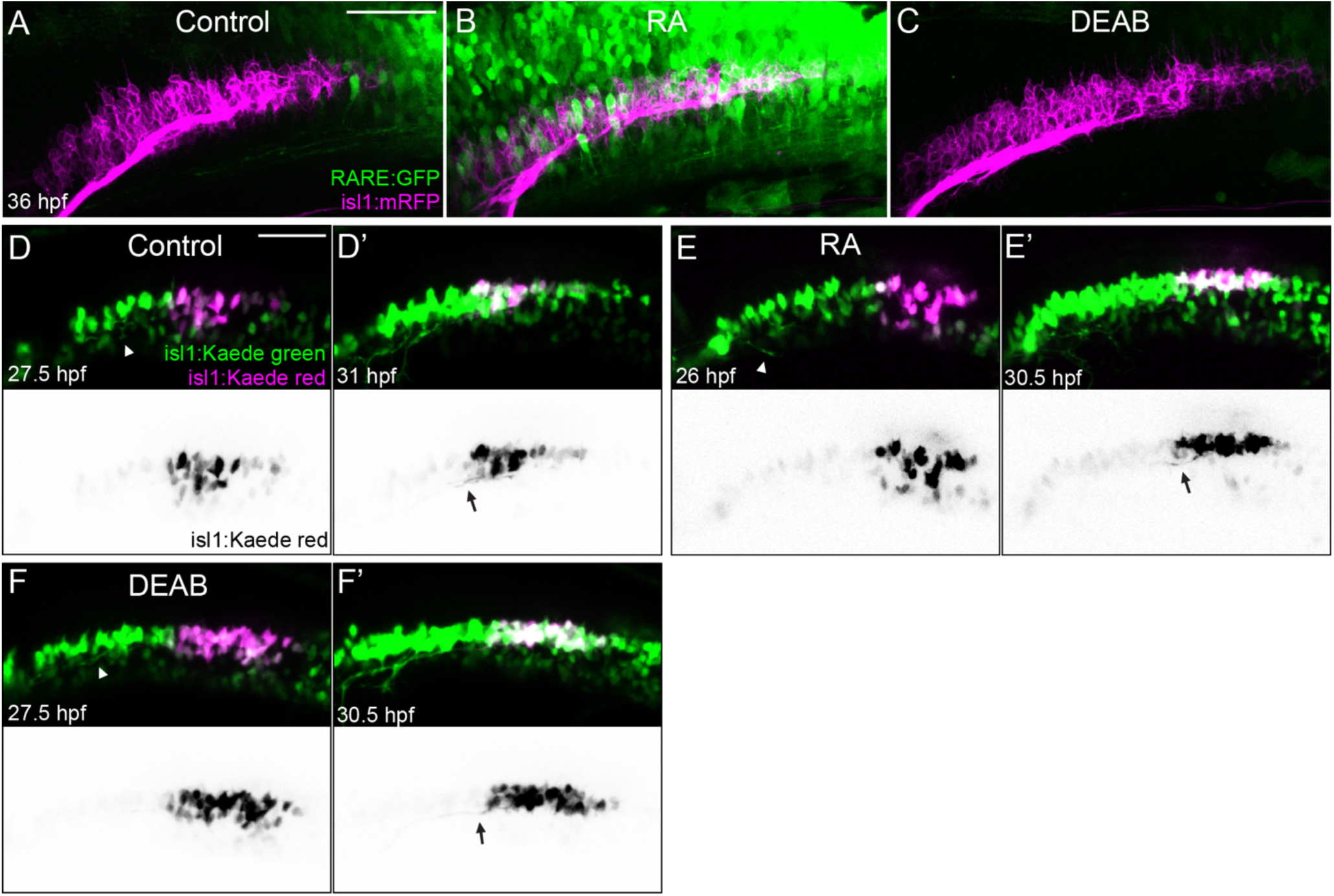
(A-C) RARE:GFP (green) levels in the vagus (magenta) territory in Control (A), RA(B), and DEAB(C) treated embryos. (D-F) Time lapse stills of Control (D), RA (E) and DEAB (F) treated embryos showing the onset of axon formation in anterior (green) and posterior (magenta) mX neurons. Black & white images are color-inverted views of the Kaede-red channel. (D, E, F) arrowheads show emerging anterior axons. (D’, E’, F’) arrows show emerging posterior axons. All images are lateral views oriented with anterior to left. Scale bars = 50 μm.

**Supplementary Figure 2, related to Figure 4.**
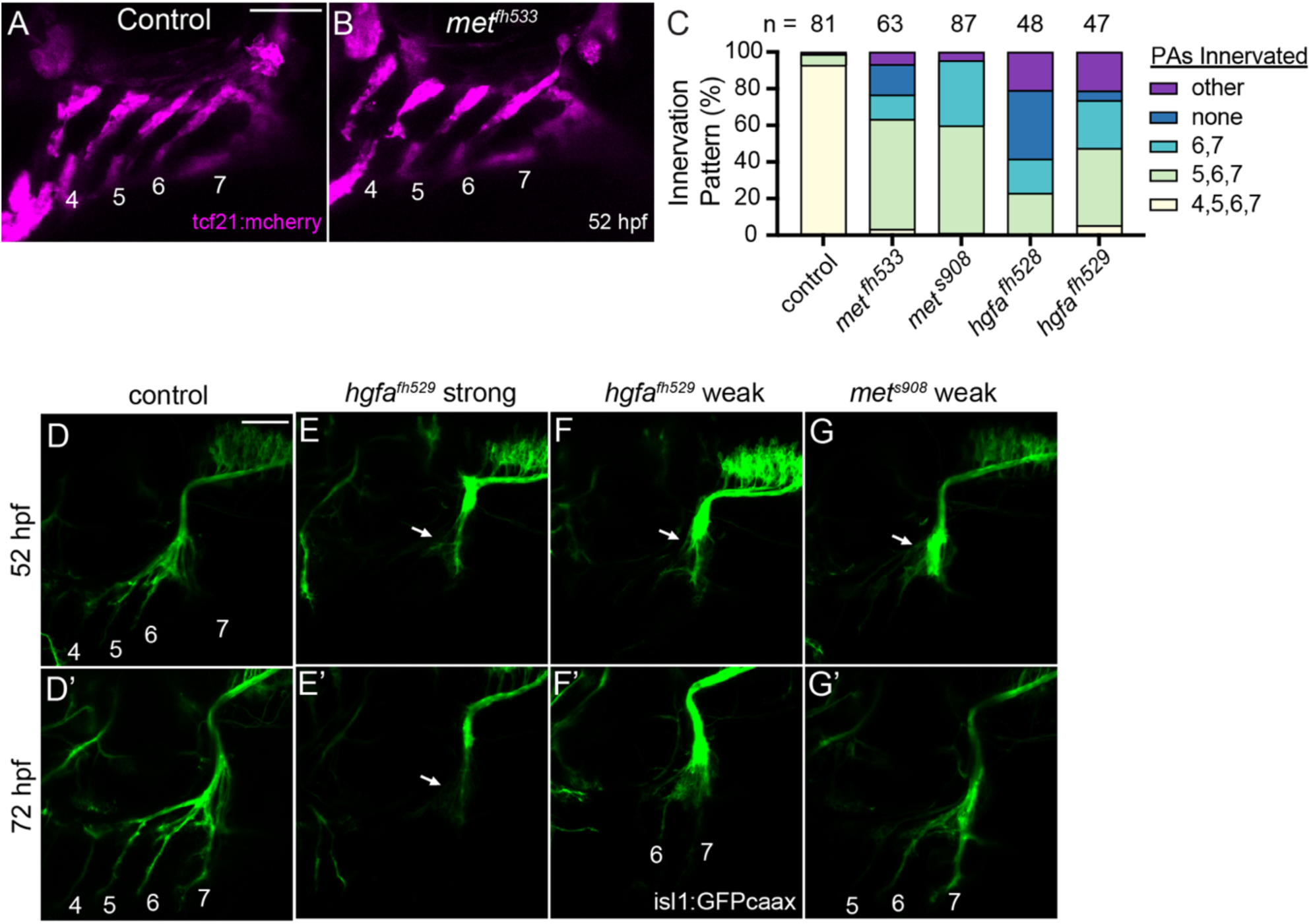
(A-B) There is no difference in the organization of the PA muscle progenitors between control (A) and *met^fh533^* (B) embryos. (C) Quantification of arch innervation pattern frequency at 72hpf in *met* and *hgf*a mutants from (D-G and Fig. 4G-K). (D-G) Paired 52hpf and 72hpf mX nerve structure in control (D), *hgfa^fh529^* (E-F), and *met^s908^* (G) embryos. Phenotypes of varying strengths are seen. At 52hpf, in all cases mX axons (green) have stalled before entering the arches (arrows) (E-G), whereas in controls axons have already innervated PAs 4, 5, and 6 (D). At 72 hpf, in the strong phenotype axons remain stalled and fail to enter the PAs (arrow) (E’), while in the weak phenotype some axons escape and variably innervate PAs 5, 6, and 7 (F’-G’). Numbers indicate innervated PAs. All images are lateral views oriented with anterior to left. Scale bars = 50 μm.

**Supplementary Figure 3, related to Figure 5.**
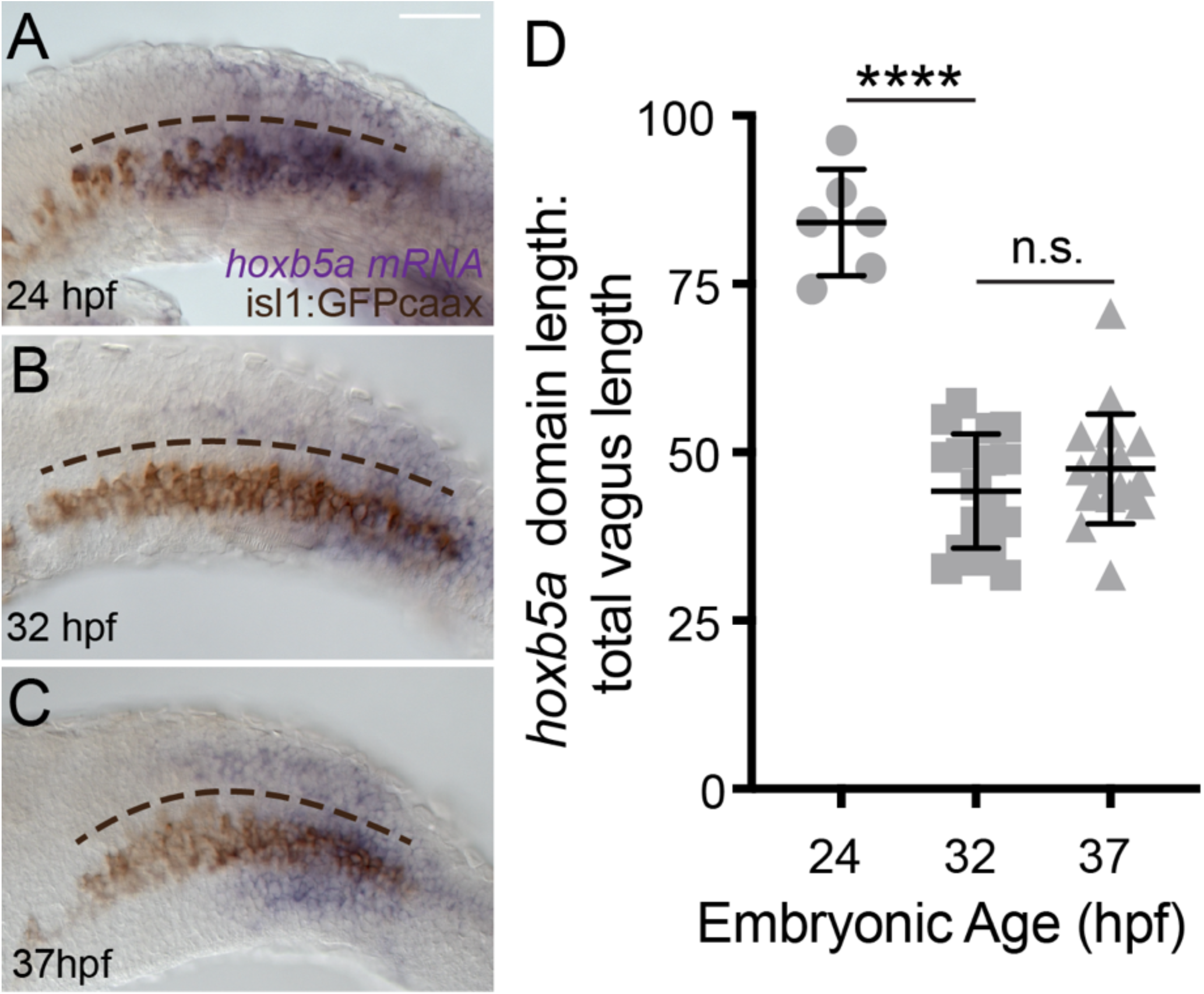
(A-C) *hoxb5a* RNA *in situ* hybridization time series showing *hoxb5a* expression receding towards the posterior over time in the mX nucleus. mRNA expression is purple and the mX neurons marked by isl1:GFP are brown. The A-P span of the mX nucleus is indicated by the curved dotted line. (D) Quantification of *hoxb5a* expression domains over time represented in (A-C). Data represent mean ± SEM. t-test ****P<0.0001, ns = not significant. All images are lateral views oriented with anterior to left. Scale bars = 50 μm.

**Supplementary Movie 1, related to Figure 4**

Time lapse of control, RA, and DEAB treated embryos, represented in (Fig S1D-F), showing the onset of axon formation in anterior (green) and posterior (magenta) mX neurons. Black & white images are color-inverted views of the Kaede-red channel.

**Supplementary Movie 2, related to Figure 4**

Time lapse of early axon targeting in control (left) and *met^s908^* mutant (right) embryos starting at 28 hpf. PA formation (magenta) and initial axon outgrowth (green) are indistinguishable between conditions, but, whereas control axons extend to the PAs, *met* mutant axons stall prior to reaching the PAs.

**Supplementary Movie 3, related to Figure 4**

Time lapse of late axon targeting in control (left) and *met^s908^* mutant (right) embryos starting at 50 hpf. PA structure (magenta) is indistinguishable between conditions but, whereas control axons fully innervate PAs 4-7, *met* mutant axons predominantly stall, with a small subset escaping to innervate PAs 5,6, and 7.

